# Combination of an inject-and-transfer system for serial femtosecond crystallography

**DOI:** 10.1101/2022.02.08.479470

**Authors:** Keondo Lee, Jihan Kim, Sangwon Baek, Jaehyun Park, Sehan Park, Jong-Lam Lee, Wan Kyun Chung, Yunje Cho, Ki Hyun Nam

**Affiliations:** Department of Mechanical Engineering, Pohang University of Science and Technology, Pohang, Republic of Korea; Department of Life Science, Pohang University of Science and Technology, Pohang, Republic of Korea; Department of Materials Science and Engineering, Pohang University of Science and Technology, Pohang, Republic of Korea; Pohang Accelerator Laboratory, Pohang University of Science and Technology, Pohang, Republic of Korea; Department of Chemical Engineering, Pohang University of Science and Technology, Pohang, Republic of Korea; POSTECH Biotech Center, Pohang University of Science and Technology, Pohang, Republic of Korea

**Keywords:** serial crystallography, sample delivery, injection, fixed-target scanning, viscous medium

## Abstract

Serial femtosecond crystallography (SFX) enables the determination of the room-temperature crystal structure of macromolecules without causing radiation damage and provides time-resolved molecular dynamics by pump-probe experiments. In the SFX experiment, the injector-based sample delivery method continuously provides fresh crystals to X-rays, and the fixed-target scanning method can be programmed to move the crystals to the desired location. This study introduces a combination of the inject-and-transfer system (BITS) method for sample delivery for SFX experiments, a hybrid injection, and a fixed-target scanning method. BITS allows solution samples to be reliably deposited on an ultraviolet ozone (UVO)-treaed polyimide films at flow rates as low as 1 nl/min. In application of BITS in SFX experiment, the lysozyme crystal samples were embedded in a viscous lard medium and injected at a 50–100 nl/min flow rate through a syringe needle onto an UVO-treated polyimide film mounted on a fixed-target scan stage. The deposited crystal sample on film were raster scanned to XFEL by motion stage in the horizontal and vertical directions. Using this method, we successfully determined the room-temperature structure of lysozyme at 2.1 Å resolution. This method can be applied to the SFX experiments.

## 1. Introduction

Serial femtosecond crystallography (SFX) enables the determination of the crystal structure at room or nearly physiological temperature without causing radiation damage [1–7]. Moreover, SFX with pump-probe experiments enables visualization of the time-resolved molecular dynamics of macromolecules [8–13]. In the SX experiment, many crystals were delivered to the X-ray interaction point and exposed to X-rays only once. Various sample delivery methods such as injectors [14–16], syringes with viscous medium [17–20], capillary methods [21, 22], fixed-target scanning [23–26], electrospinning [27], and microfluidic devices [28–30] have been applied to deliver a large number of crystals into X-ray interactions in a serial manner.

Among them, the liquid jet injector has the advantage of maintaining the crystal hydrate environment and providing fresh crystals from the injector to X-rays with reduced exposure to the atmosphere [14]. However, a high flow rate is generally required to provide a stable injection flow. Therefore, it is difficult to apply liquid beam injectors to XFEL with a low repetition rate in terms of sample consumption. Moreover, due to the XFEL source’s pulsed nature, large amounts of crystals are not transmitted to X-rays [31]. Recently, to reduce sample consumption in the injector method, a segmented flow generator method wherein droplets containing crystals are segmented with immiscible oil in a microfluidic device has been developed [31].

The method of delivering the crystals embedded in a viscous medium via an injector or syringe provides a stable injection stream even at a low flow rate and is therefore widely applied in XFEL facilities with low repetition rates or synchrotrons [15, 17, 32]. Several types of viscous media, such as lipidic cubic phase (LCP) and hydrophobic and hydrophilic injection matrices, have been developed and successfully applied to SX experiments [20]. However, depending on the crystal characteristics and the crystallization solution composition, the viscosity of the viscous medium may change, which may produce an unstable injection stream [20]. In such cases, an effort is required to find a stable injection condition, which lowers the efficiency of data collection and increases the sample consumption during the optimization of the injection stream. To solve this problem, the delivery of crystals embedded in a viscous medium via capillary or single-channel microfluidics aligned to the X-ray interaction point was developed [22, 30]. Since the inner channel of the capillary or microfluidics was aligned to the X-ray beam path, all crystals continually pass through the inner channel of those devices and are exposed to the X-ray. Although these methods have been successfully proven in serial synchrotron crystallography (SSX) experiments, glass or polyimide channels may be physically damaged when exposed to intense XFEL.

The fixed target scanning method has advantages in that it consumes fewer crystal samples compared to the sample delivery method using an injector, minimizes physical impact on the crystal during X-ray delivery, and can be programmed and delivered to the desired location [23, 25, 26, 33–35]. Although radiation damage has not yet been reported in the fixed target scanning method, theoretically, radicals are generated after XFEL is passed [26], so if they are diffused, they may affect the diffraction data of later collected crystals. In such cases, it may be limited to time-resolved studies using an optical laser with caged molecules.

In contrast, a hybrid-type sample delivery method that combines injection and motion was developed. In the mix-and-diffuse [36] or drop-on-drop [37] method, crystal samples are placed on the polyimide tape by an injector, and these crystals are delivered to the X-ray interaction point by driving the conveyor belt. This method enables liquid-based time-resolving studies for various time delays by controlling the mixing speed and speed of the conveyor belt [36, 37]. In the liquid application method for time-resolved analyses (LAMA) [38] approach, the crystals are injected into a fixed-target chip, and these crystals are delivered to the X-ray interaction point by the translation stage motion. This method enables a time-resolved SX experiment using an optical laser or liquid application. These hybrid sample delivery systems are useful for application in time-resolved SFX experiments, but they are physically difficult to apply to our sample environment.

In this study, we introduce a new sample delivery method named ‘com**B**ination of **I**nject-and-**T**ransfer **S**ystem’ (BITS), which combines the advantage of delivering a fresh sample every time from the previously reported injection method and the crystal sample programming method to the desired location in the XFEL in the SFX fixed target scanning method. Lysozyme crystals were embedded in lard and extruded on a UVO-treated polyimide film through a syringe. Crystal samples were scanned by the translational motion stage in the horizontal and vertical directions. Using this method, we successfully determined the room-temperature structure of lysozyme. This sample delivery method can be applied to the SFX data collection.

## 2. Methods and materials

### 2.1. Crystallization

Chicken egg white lysozyme was purchased from Sigma-Aldrich (Cat. No. L6876, St. Louis, MO, USA). Crystal samples were obtained using the microtube batch method, as previously described [39]. Briefly, lysozyme was dissolved in a solution containing 10 mM Tris-HCl, pH 8.0, and 200 mM NaCl. Then, the lysozyme solution (50□mg/ml) was mixed with an equal volume of crystallization solution containing 0.1□M sodium acetate, pH 4.0, 5% (w/v) PEG 8000, and 3.5 M NaCl in a 1.5 ml microtube, followed by incubation at room temperature. The cubic-shaped lysozyme crystals grew to dimensions of approximately 10 × 10 × 10 □μm in a hours.

### 2.2. Crystal embedding in lard

The lard injection matrix viscous medium was prepared as previously reported [40]. The lysozyme crystals and lard were mechanically mixed in a dual syringe setup, as previously reported [20, 40]. Briefly, the lysozyme crystal suspension was transferred into a syringe. Then, the lard was dissolved by immersing in hot water (approximately 80 °C) for 10 min, transferred into a syringe, and left to stand until it solidified. The syringes containing the lysozyme crystals (200 μL) and lard (200 μL) were connected using a coupler and mixed by moving the plunger back and forth more than 30 times. Finally, the crystals embedded in the lard were transferred to a syringe connecting a needle with an inner diameter of 150 μm.

### 2.3. Ultraviolet oxygen treatment

Polyimide film (50 μm) was purchased from Suzhou Kying Industrial Materials Co.,Ltd (Suzhou, China). The UV/ozone (UVO) treatment was conducted on a 70 mm x 70 mm polyimide film with a UVO Cleaner (Ahtech LTS Co., Ltd., Anyang, Republic of Korea) for > 10 min. UVO-treated polyimide film was immediately packed under vacuum sealing to avoid the recovery of the polyimide film surface.

### 2.4. Data collection

SFX data collection was performed at the nano-crystallography and coherent imaging (NCI) experimental hutch in the Pohang Accelerator Laboratory X-ray Free Electron Laser (PAL-XFEL) [41, 42]. The photon energy and flux used in this experiment were 9500 eV and 2-5 × 10^11^ photons/pulse, respectively. The X-ray beam size was 2 μm (vertical) × 3 μm (horizontal) (full-width half-maximum) focused on using a Kirkpatrick-Baez mirror [43]. A 1000 μl syringe (BD Plastics Ltd, Sunderland, United Kingdom) containing the crystals embedded in the lard injection matrix was mounted on a Fusion 200 Touch syringe pump (Chemyx Inc., Houston, TX, USA). The sample was transferred from the syringe to the needle with an inner diameter of 790 μm along the PTFE tubing (Precigenome LLC, San Jose, CA, USA). The position of the needle was moved using an X-Y manual motion stage, and the needle tip was tilted at an angle of 30–60 ° to contact the lower-left corner of the UVO-treated polyimide film. Crystal samples were extruded at a 50-100 nl/min flow rate with a Fusion 200 Touch syringe pump (Chemyx Inc., Houston, TX, USA). The scanning stage moved from left to right without synchronization with the arrival of an FEL pulse, and then moved again to the left and then the right in a zigzag manner in the scanning direction. When moving in the vertical direction, it moved from bottom to top at intervals of 50 μm three times, moved to the right at intervals of 50 μm, then moved downward at intervals of 50 μm down three times at intervals of 50 μm, and then moved to the right again at intervals of 50 μm. This type of raster scanning motion was repeatedly performed to collect the complete three-dimensional structure data. Diffraction data were collected in a helium atmosphere at 23–25 °C using an MX225-HS detector (Rayonix, LLC, Evanston, IL, USA) with a 30 Hz readout.

### 2.5. Structure determination

The diffraction patterns image was filtered using the *Cheetah* program [44]. Hit images were indexed and scaled using *CrystFEL* [45]. The electron density map was obtained by molecular replacement with the *MOLREP* program [46] with the lysozyme crystal structures (PDB code 7WKR) as the search mode. The model structure was rebuilt using the *COOT* [47]. The final model was refined using *the PHENIX* [48]. The geometry of the final model structures was verified using *MolProbity* [49]. Structural figures were generated using *PyMOL* (https://pymol.org/). Data collection and refinement statistics are summarized in Table 1. Coordinates and structure factors have been deposited under accession code 7WUC in the Protein Data Bank.

**Table 1.** Data collection and refinement

| Data collection |  |
| --- | --- |
| Diffraction source | NCI beamline, PAL-XFEL |
| Wavelength (Å) | 1.305 |
| Temperature (K) | 296.15- 298.15 |
| Photons/pulse | $\sim 5 \times 10^{11}$ |
| Detector | MX225-HS |
| Crystal-detector distance (mm) | 123 |
| No. collected images | 85901 |
| No. of hits | 27413 |
| No. of indexed patterns | 22303 |
| Space group | $P4_32_12$ |
| $a, b, c$ (Å) | 79.48, 79.48, 38.48 |
| $\alpha, \beta, \gamma$ (°) | 90, 90, 90 |
| Resolution range (Å) | 80.0 – 2.10 (2.17 – 2.10) |
| No. of unique reflections | 7521 (724) |
| Completeness (%) | 100 (100) |
| Redundancy | 590.7 (377.5) |
| SNR | 7.38 (1.57) |
| *CC* | 0.9866 (0.4766) |
| *CC* * | 0.9966 (0.8034) |
| *R*<sub>split</sub> (%) <sup>a</sup> | 10.44 (71.03) |
| Overall *B* factor from Wilson plot (Å<sup>2</sup>) |  |
| **Refinement** |  |
| Resolution (Å) | 39.74–2.10 (2.26–2.10) |
| σ cutoff | $F > 1.35\sigma(F)$ |
| Final *R*<sub>work</sub> <sup>b</sup> | 0.220 (0.3563) |
| Final *R*<sub>free</sub> <sup>c</sup> | 0.249 (0.3833) |
| B-factor (Averaged) |  |
| Protein | 43.87 |
| Water | 46.45 |
| R.m.s. deviations |  |
| Bond lengths (Å) | 0.003 |
| Bond angles (°) | 0.664 |
| Ramachandran plot (%) |  |
| favoured | 97.64 |
| allowed | 2.36 |

<sup>a</sup> $R_{split} = \left(1/\sqrt{2}\right) \cdot \frac{\sum_{hkl} |I_{hkl}^{even} - I_{hkl}^{odd}|}{\frac{1}{2} \sum_{hkl} |I_{hkl}^{even} + I_{hkl}^{odd}|}$
<sup>b</sup> $R_{work} = \sum ||F_{obs}| - |F_{calc}|| / \sum |F_{obs}|$ , where $F_{obs}$ and $F_{calc}$ are the observed and calculated structure-factor amplitudes respectively.
<sup>c</sup> $R_{free}$ was calculated as $R_{work}$ using a randomly selected subset (9.47%) of unique reflections not used for structure refinement.

## 3. Results and Discussions

### 3.1. Combination of inject-and-transfer system

We developed the comBination of Inject-and-Transfer System (BITS) from two perspectives; (i) it was intended to be used immediately without effort to create a stable injection stream when the injection stream, including crystals delivered from the injector or syringe, is provided unstable. (ii) We tentatively aimed to develop a novel mix-and-injection method for time-resolved SFX. To achieve these requirements, we first considered sample delivery using microfluidic chips. However, when the XFEL exposed the microfluidics chip consisting of polydimethylsiloxane (PDMS) or polyimide, a hole in the chip was created due to radiation damage. In addition, the sample path was changed by the bubble generated when the XFEL penetrated the solution, and the channel clogging problem also occurred in the channel due to crystal accumulation. Therefore, we concluded that crystal samples should be exposed to the atmosphere to avoid these radiation damage issues when exposed to XFEL. Regarding this aspect, the mix-and-inject injector is attractive [50, 51]; however, it requires a fast flow rate to create a stable injection stream. Therefore, it is not suitable because the consumption of crystal samples is overwhelming when applied in PAL-XFEL with a low repetition rate. Then, the mix-and-diffuse [36] or drop-on-drop [37] method using the conveyor belt approach and the LAMA [38] approach were also considered suitable systems for our research goals. However, this instrument may require considerable effort for precise alignment to pass the XFEL to the sample, so it is difficult to apply when obtaining very short beamtime. In addition, to apply these devices to our sample environment, there was a problem wherein the sample environment, including the sample chamber, had to be reconfigured.

Therefore, we basically benchmarked the previous hybrid sample delivery method and developed a new BITS (comBination of Inject-and-Transfer System) sample delivery method that considered our sample delivery environment. The BITS consisted of injection and motion systems, and the sample chamber and motion stage utilized the previously developed FT-SFX chamber system. The main components of BITS can be divided into injection, which delivers the crystal sample from the syringe and tube to the film via a syringe needle, and a motion stage, which moves the crystal sample from the film to the point of X-ray interaction (**Fig. 1**). The syringe needle tip, where the crystals are ejected, is adjusted with a manual stage to attach the film where the crystals are placed. The crystal sample forms a small drop at the needle tip, and when they come into contact with the film, they are deposited on the film according to the flow rate. Meanwhile, when the needle tip does not directly contact the PI film, a sample drop is formed at the end of the needle tip by surface tension, and as the drop grows, data collection cannot be performed until it reaches the polyimide film. This causes an increase in the amount of unnecessary sample consumption and impairs data collection efficiency. Accordingly, the needle tip must contact the polyimide film during device installation as well as during the data acquisition. Therefore, it is important that the polyimide film is installed flat on the frame and that the translation stage is accurately aligned horizontally and vertically with the needle. When the injected crystal sample was placed on the film, the movement of the translation stage transferred the sample to the X-ray position and exposed it to the XFEL.

**Figure 1.**
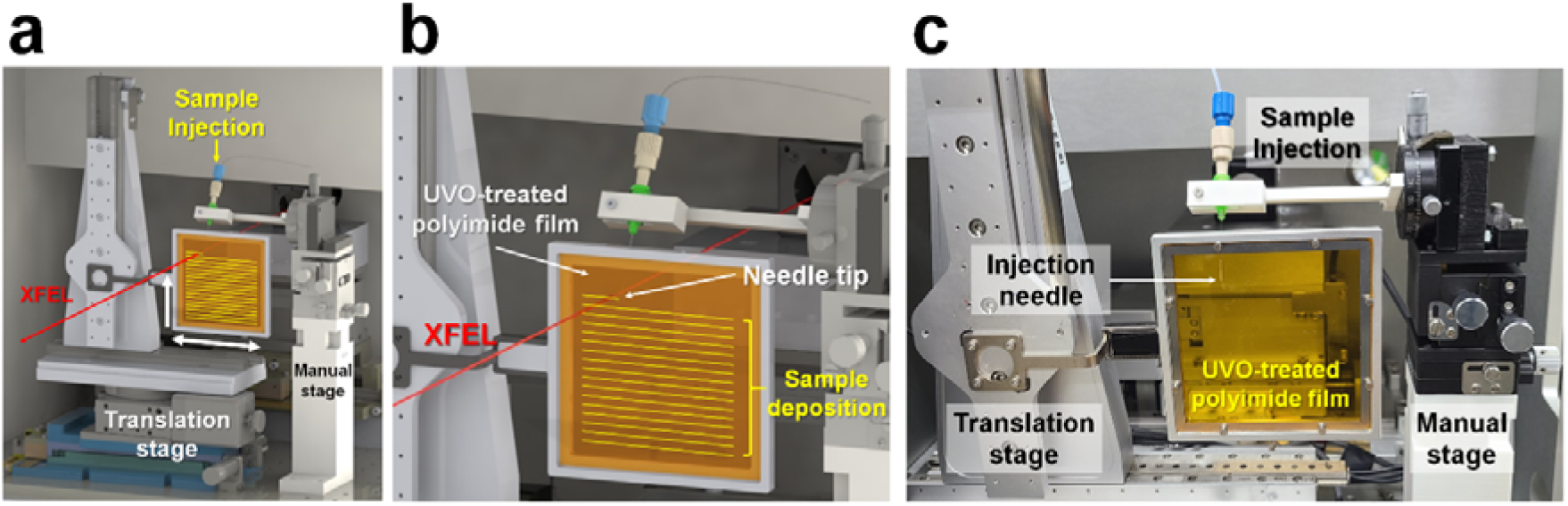
Schematics of the experimental setup of the BITS. (a) BITS is composed of the injection and translation system. (b) The protein crystal sample (yellow) is injected from the needle and deposited on the UVO-treated polyimide film, the needle position is fixed, and the sample position is moved to the XFEL path (red line) by the motion stage. (c) Photo of experimental setup of the BITS.

Collectively, the BITS method is a combination of the conventional injection method and fixed target scanning method. It has both the advantages of an injector that always delivers a fresh sample and a fixed target scanning method that allows the sample to be freely programmed to the desired location. In addition, because the stability of the injection flow is not critical, it can be delivered at very low flow rates, reducing sample consumption.

### 3.2. Sample loading

To demonstrate the stable sample loading to the film, an initial injection experiment on polyimide film was performed using a crystallization solution containing 0.1 M sodium acetate, pH 4.0, 5% (w/v) PEG8000, and 3.5 M NaCl. The results show that rather than uniformly depositing the crystal solution on the polyimide film, a drop was formed at the syringe needle tip, and it often settled on the polyimide film with a random-sized drop (**Fig. 2a**). This phenomenon favors the droplet formation at the needle tip rather than the deposition of the crystallization solution on the film because the polyimide film surface is very hydrophobic [52]. Moreover, it was considered that the crystallization solution prefers to maintain the drop formation on the polyimide film because the crystallization solution has no affinity for the polyimide film hydrophobic surface, even if the droplet becomes large and settles on the film.

**Figure 2.**
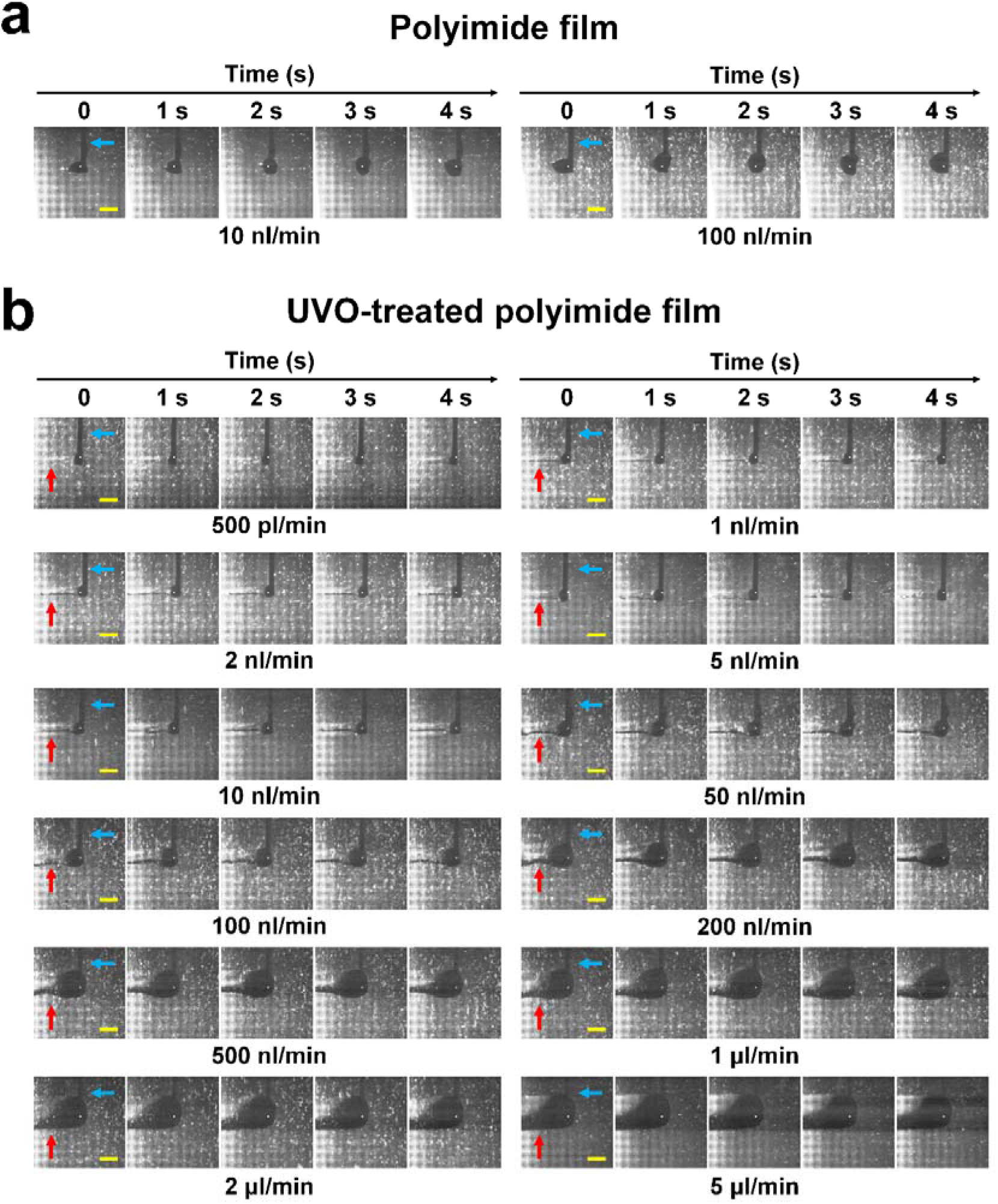
Sample injection experiment on polyimide film. (a) Snapshots of the sample injection on polyimide film at flow rates of 10 and 100 nl/min. (b) Snapshots of the sample injection on UVO-treated polyimide film at flow rates of 0.5, 1, 2, 5, 10, 50, 100, 200, 500, 1000, 2000 and 5000 nl/min.

To solve this problem, the hydrophilicity of the polyimide film surface was increased by UV/ozone (UVO) surface pre-treatment. The results showed that the crystallization solution was effectively deposited on the UVO-treated polyimide film (**Fig. 2b**). Therefore, the polyimide film hydrophilic surface using UVO-treatment helped the sample settle on the film and was suitable for application in the BITS. Meanwhile, the hydrophilic surface of the UVO-treated polyimide film can convert to its original hydrophobic surface over time. Therefore, applying the polyimide film immediately after UVO treatment would be ideal.

The stability of the injected sample and sample consumption varies depending on the injection flow rate and translation stage speed. Therefore, the target scanning was collected at 30 Hz at 50 μm intervals in the horizontal direction, and the crystallization solution was injected while moving at the translation speed of 1.5 mm/s. Then, the sample pattern on the UVO-treated polyimide film according to various injection flow rates was analyzed (**Fig. 2b**). As a result, the crystallization solution was stably mounted on the UVO-treated PI film at an injection rate of 1-100 nl/min. At 500 pl/min, it was determined that the sample was continuously injected on the UVO polyimide film; however, in this case, it was expected that a dehydration problem could occur experimentally. The sample was uniformly deposited on the film even at a flow rate of > 200 nl/min, but the width and thickness of the sample were unnecessarily wide or thick, and the solution that was not deposited on the film formed droplets on the needle tip. This is an excessive injection flow rate, which temporarily causes unnecessary crystal consumption.

On the other hand, we also used a nylon mesh as the crystal settling plane material and stably and uniformly mounted the crystal solution. However, the needle tip position was often slightly changed while the translation stage was moving (Fig. S4). This is caused by a physical effect that occurs when the needle tip is in contact with a nylon mesh with an uneven surface and moves. Nevertheless, it is possible to collect data by injecting crystal samples on nylon mesh, but a UVO-treatment polyimide film was used in this study to deliver samples in an optimized state.

### 3.3. XFEL data collection

To demonstrate the BITS application, we performed an SFX experiment using lysozyme as a model sample. In the initial XFEL data collection, the lysozyme crystal suspension was loaded onto a UVO-treated polyimide film through a syringe needle to expose the X-rays. During SFX data collection, lysozyme diffraction data could be collected for several minutes; however, the crystal diffraction pattern decreased as the injection time increased, and the solution scattering was overwhelmingly increased. When the sample tubing and syringe needle were disassembled, we confirmed that the crystal sample was clogged at the needle inlet. We considered that when the crystal suspension was delivered at a low flow rate, the crystals sank to the tube and needle by gravity, and a large number of crystals clogged the needle inlet, with only the crystallization solution flowing after clogging. We suggested that it may be possible to reduce the crystal clogging by gentle shaking using an anti-settler where the crystals are stored or by minimizing the distance of the tubing through which the crystals pass.

In contrast, our previous study investigated how the delivery of crystals mixed with viscous media such as agarose, gelatin, or shortening prevented the crystals from sinking in gravity without clogging in capillary and single-channel microfluidics [22, 30]. Similarly, lysozyme crystals were embedded in the lard material in this experiment to prevent clogging of the crystals in the needle and tubing. Lard was used to produce a stable injection stream at a low flow rate in the previous SX experiment, and a stable stream was produced when the lard content was > 80% in the mixture of lard and protein suspension [40]. In this experiment, the crystal suspension and lard ratio was 5:5; therefore, an injection stream was not created, and it was used only to prevent crystals from sinking. As a result, crystals embedded in lard were stably delivered to UVO-treated polyimide films without clogging in tubing or syringe needles. The lard used in this study showed background scattering at about 4.3 Å, which had no effect on structural determination, however, if the diffraction intensity of the target crystal was weak or to minimize background noise, viscous materials with low background scattering should be used. On the contrary, in this experiment, a mixture of viscous material and crystals, which cannot create an injection stream, was injected into the film. However, data collection was possible by directly utilizing the injection stream on the film, similar to general sample delivery using a viscous medium in the SX experiment. In contrast, to prevent the crystal from sinking by gravity using a viscous material, it is necessary to select a viscous material that does not have a physical and chemical effect during the mixing and storage of the crystal and the viscous material.

Our injection test showed that the injection sample was stably mounted on the film, even at 0.5 nl/min (**Fig. 2b**). However, the crystallization solution of lysozyme contains a 3.5 M NaCl concentration, and prolonged exposure to the atmosphere during data collection may cause salt crystal growth and changes in the crystal lattice by dehydration. Therefore, to prevent this in advance, crystals embedded in the lard were injected at 50–100 nl/min in the actual experiment. In BITS, if there were no dehydration issues of the crystallization solution, the crystal sample would be ejected at a lower flow rate.

The width of the injected crystals embedded in the lard on the UVO-treated polyimide film was approximately 300 μm (**Fig. 3a**). Because the size of the beam was within 5 μm (FWHM), most crystal samples were not exposed to X-rays when translation solely moved in the horizontal direction during data collection (**Fig. 3b**).

**Figure 3.**
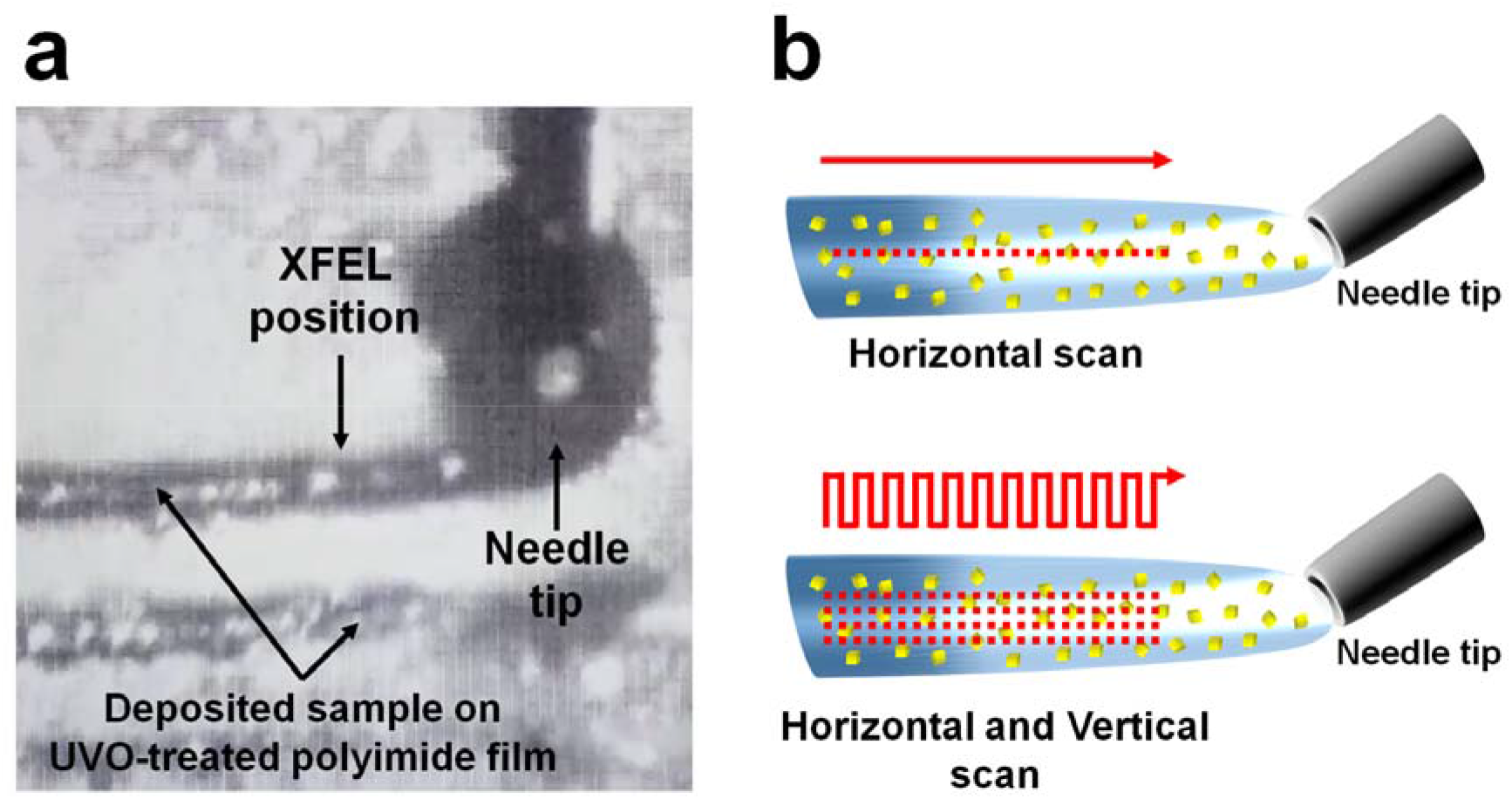
Injection and data collection strategies in the BITS approach. (a) Snapshot of the injection of crystals embedded in lard on UVO-treated polyimide film for data collection using BITS. (d) Data collection strategy in BITS. (up) When the motion stage moves in one direction, X-rays are only exposed to one position in the injected sample, and most crystal samples are not exposed to X-rays. (down) In BITS, more data can be collected from the injected sample because the motion stage moves both horizontally and vertically.

Alternatively, BITS can be programmed to move the sample to the desired location. Therefore, the width of the injected sample is not an important factor in terms of sample consumption. In this experiment, to avoid radiation damage to the surrounding sample when XFEL was transmitted, crystals were scanned at 50 μm intervals. During data collection, the process of scanning the injected sample vertically at 50 μm intervals for three scan points and moving it 50 μm to the right was repeated. The applied injection sample shows various injection sample widths on the film depending on the used inner needle (or nozzle) diameter and the crystallization solution viscosity when applying BITS. However, the injected sample width is not critical, as data collection can be accomplished by simply programming the scan interval according to the injected sample width. Conversely, in the existing injector/syringe, X-rays can be exposed only at one point at the injection stream, whereas in BITS, both can be exposed to X-rays vertically and horizontally at a lower flow rate so that sample consumption can be significantly reduced.

XFEL data were collected using BITS with lysozyme crystals embedded in lard material. ). A total of 85,901 images were collected for 50 min, and the hit images containing the Braggs peaks and indexed images were 27,413 (hit rate: 31.91%) and 22,302 (indexing rate: 81.35%), respectively. The data were processed up to 2.1 Å, and overall completeness, signal-to-noise ratio (SNR), R_split_, Pearson correlation coefficient (CC), and CC* were 100, 7.38, 10.44, 0.9866 and 0.9966, respectively (**Table 1**). The R_work_ and R_free_ of the room temperature lysozyme structure were 0.220 and 0.249, respectively (**Table 1**). Lysozyme showed a clear electron density map for Lys19-Leu147 (**Fig. 4a**). No significant radiation damage was observed in the disulfide bonds (Cys24-Cys145, Cys48-Cys133, Cys82-Cys97, and Cys94-Cys112) in lysozyme (**Fig. 4b**). The room-temperature structure of lysozyme by BITS showed high similarity with the previously reported room-temperature structure of lysozyme delivered in polyacrylamide (PDB code: 6IG6) [39], shortening (6KCB)[53], lard (7CJZ) [40], and beef tallow (7E02) [54], with an RMSD of 0.111-0.128 Å.

**Figure 4.**
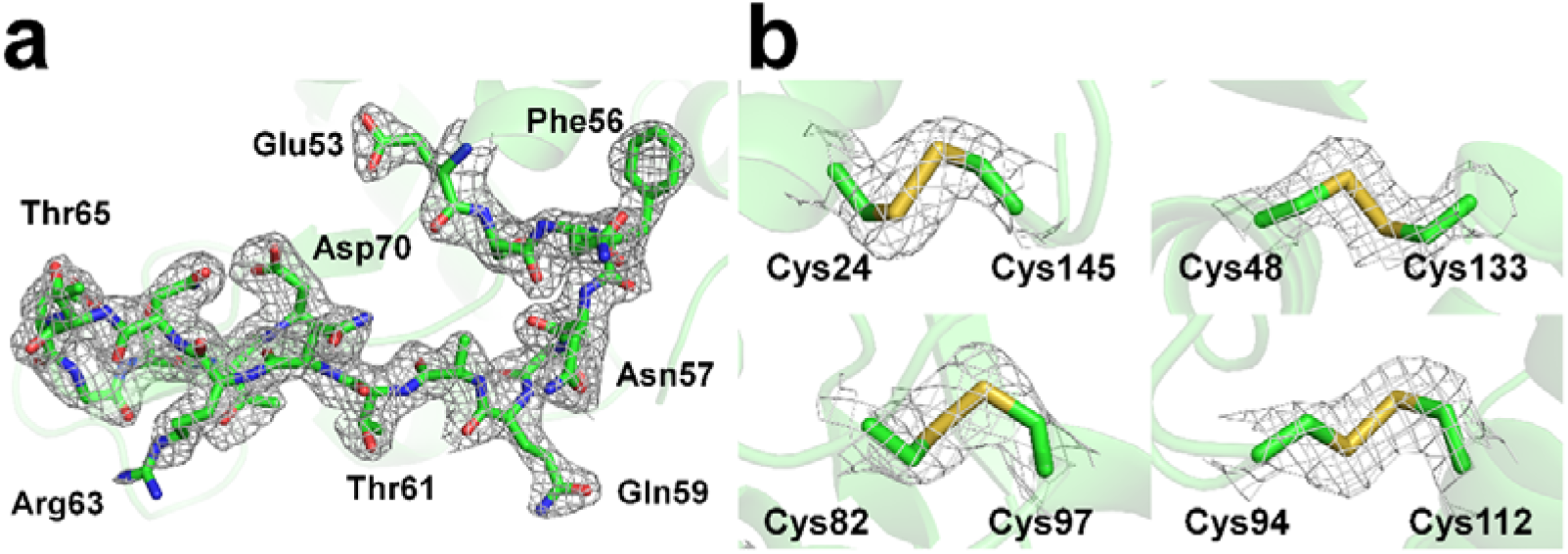
Electron density map of room-temperature lysozyme embedded in lard delivered by BITS. (a) 2 mFo-DFc electron density map (grey mesh, 1.0σ) of lysozyme. (b) 2 mFo-DFc (grey mesh, 1.0σ) and mFo-DFc (green, 3σ; red,□-□3σ) electron density maps around the disulfide bonds of lysozyme.

## 4. Conclusion

In this study, we introduced the BITS, a hybrid injection method, and a fixed-target scanning method. Using this approach, we successfully collected XFEL diffraction data and determined the lysozyme room-temperature structure. This method does not require any effort to create a stable injection stream in the existing injector/syringe-based sample delivery method, thus delivering samples at very low flow rates. In addition, even if the width of the injection sample applied to the experiment varied, most of the injected samples could be scanned by programming the motion stage. As a result, compared to the sample delivery method using the existing injector/syringe, it requires less effort for sample injection and has the advantage of significantly reducing sample consumption. In future studies, it will be possible to conduct time-resolving studies of mix-and-diffuse or drop-on-drop methods on BITS using the previously reported T-junction or two injectors’ concepts with piezoelectric control. Furthermore, our method can be applied to the SFX data collection.

## Acknowledgements

We thank the beamline staff at NCI bemline at Pohang Accelerator Laboratory X-ray Free Electron Laser (PAL-XFEL) for their assistance with data collection. The XFEL experiments were carried out at the NCI endstation at PAL-XFEL (Proposal Nos: 2021-2nd-NCI-013). The authors thank Global Science experimental Data hub Center (GSDC) at Korea Institute of Science and Technology Information (KISTI) for computational support. This work was supported by the National Research Foundation of Korea (NRF-2017M3A9F6029736; NRF-2020M3H1A1075314; NRF-2021R1I1A1A01050838).

